# Reducing High Pressure Processing Costs: Efficacious Alternatives to Current Standard Procedures in the Food Manufacturing Industry

**DOI:** 10.1101/2022.12.15.520669

**Authors:** Anita Scales Akwu, Abimbola Allison, Jayashan Adhikari, Monica Henry, Wendelyn Inman

**Affiliations:** Tennessee State University, College of Agriculture, Human, and Natural Sciences, Public Health, Health Administration, & Health Sciences, Nashville, TN 37203

## Abstract

As a result of recent advancements in design and optimization of high-pressure processing units, the technology is gaining rapid adoption across various sectors of food manufacturing, thus requiring extensive microbiological hurdle validation studies for efficacious and feasible utilization of the technology. Commercial adoption of high-pressure processing is gaining momentum of industrial importance because of recent advances in the engineering of pressure-based pasteurization units. With tremendous ability of plethora of microorganisms to move towards fitness through vertical and horizontal gene transfer mechanisms, prevention of natural and anthropogenic pathogens of public health concern is a daunting task and a moving target. Current study discusses, Reducing the Cost Associated with High Pressure Processing: Efficacious Alternatives to the Current Standard Procedure in the Food Manufacturing Industry, with microbiological challenge studies for inactivation of the pathogen exposed to various times and intensity levels of elevated hydrostatic pressure (Pressure BioScience Inc.).

Elevated hydrostatic pressure is a non-thermal procedure that exposes pathogens to pressures of up to 80,000 PSI (>550 MPa). Various times (3, 4, and 5 minutes) at pressure intensity levels of 600 MPa, (87K PSI), 550 MPa (79K PSI), 480 MPa (70K PSI), 415 MPa (60K PSI), and 345 MPa (50K PSI) of elevated hydrostatic pressure (Hub880 Explorer, Pressure BioScience Inc), were investigated at 4°C and for 45°C for inactivation of Shiga toxin-producing *Escherichia coli* O157:H7 (STEC) (ATCC numbers BAA 460, 43888, 43894, 35150, 43889, and 43890) respectively, ‘Big Six’ non-O157 Shiga toxin-producing *E. coli* (nSTEC) (ATCC numbers BAA 2196, 2193, 2215, 2440, 2219, and 2192) respectively, *Salmonella* serovars (ATCC numbers 13076, 8387, 6962, 9270, and 14028) respectively, and *Listeria monocytogenes* (ATCC numbers 51772, 51779, BAA 2657, and BAA 13932). Studies were conducted in two biologically independent repetitions as a blocking factors of a randomized complete block design containing three repetitions per time/treatment within each block, analyzed statistically using GLM procedures of SAS 9.4 software at type one error level at 5% using Tukey- and Dunnett-adjusted ANOVA. A Barocycler Hub840 unit (Pressure BioScience Inc., Southeastern, MA), equipped with a water jacket and circulating water bath for precise application of hydrostatic pressure and controlled temperature was utilized.

Up to 0.95 and 2.60 log reductions (P<0.05) of non-habituated Shiga toxin-producing *Escherichia coli* Non-O157 at planktonic stages were achieved using application of pressure at 345 MPa and 550 MPa for 5 minutes and 4 minutes, respectively at 4°C. Up to 4.42 and 5.10 log reductions (P<0.05) of non-habituated Shiga toxin-producing *Escherichia coli* Non-O157 at planktonic stages were achieved using application of pressure at 345 MPa and 480 MPa for 5 minutes and 4 minutes, respectively at 45°C. Up to 1.63 and 3.14 log reductions (P < 0.05) of non-habituated *Listeria monocytogenes* at planktonic stages were achieved using application of pressure at 345 MPa and 600 MPa for 5 minutes and 3 minutes, respectively at 4°C. Up to 4.91 and 6.37 log reductions (P < 0.05) of non-habituated *Listeria monocytogenes* at planktonic stages were achieved using application of pressure at 550 MPa and 480 MPa for 4 minutes, respectively at 45°C. Up to 2.87 and 5.82 log reductions (P< 0.05) of non-habituated *Salmonella* serovars at planktonic stages were achieved using application of pressure at 345 MPa and 550 MPa for 5 minutes and 4 minutes, respectively at 4°C. Up to 5.17 and 6.79 log reductions (P< 0.05) of non-habituated *Salmonella* serovars at planktonic stages were achieved using application of pressure at 415 MPa and 600 MPa for 5 minutes and 3 minutes, respectively at 45°C. Up to 0.86 and 1.35 log reductions (P<0.05) of non-habituated Shiga toxin-producing *Escherichia coli* O157:H7 at planktonic stages were achieved using application of pressure at 345 MPa and 480 MPa for 5 minutes and 4 minutes, respectively at 4°C. Up to 2.02 and 6.12 log reductions (P<0.05) of non-habituated Shiga toxin-producing *Escherichia coli* O157:H7 at planktonic stages were achieved using application of pressure at 345 MPa and 550 MPa for 5 minutes and 4 minutes, respectively at 45°C. Results of this study could be incorporated as a part of predictive public health microbiology modeling and risk assessment analysis for prevention of pathogen related disease and illness episodes.

## 1. Introduction

Consumers are very particular about the food that they consume, and many factors are considered. Some factors include freshness, natural and absent of preservatives and synthetic additives (Arora and Chauhan, 2019). Additionally, food safety and quality are very important to the consumers, but without compromising nutritional and sensory characteristics of the food products. Desired food products by consumers are those that maintain the original texture, color, and nutrients, while also being free of microbiological spoiling microflora. Though customers desire many things in their food products, pressure treated products are slightly more expensive than traditional commodities.

High Hydrostatic Pressure Processing (HPP) is constantly evolving and expanding its capabilities and venues. HPP started out as a very basic model and has been consistently being modified and enhanced. HPP utilizes water as its medium as it transfers water between pressures ranging from 100 through 1000 MPa in a confined vessel (Arora and Chauhan, 2019); in which the outcomes are food products with bacterial devaluation and elongated shelf-life (Nath et al., 2019). There were a total of five major components that collectively form the HPP unit; which include a pressure vessel, two end closures, yoke, a pressure-creating device, and instrumentation and controls (pressure measurement, temperature measurement, flow measurement, and level measurement) (Arora and Chauhan, 2019). HPP appeases inimical pathogenic and vegetative spoilage bacterium alongside of specifically chosen catalysts (Arora and Chauhan, 2019). The HPP processing temperatures range from 0 to 100 °C (Arora and Chauhan, 2019). *Ohara et al*., *2015* discussed that some food products functional characteristics are enhanced from the process of HPP. Gamma-aminobutyric acid in brown rice, buck wheat, soybean, Vitamin C content in potatoes and avocados, and the lycopene content in tomatoes were all specifically mentioned. High pressure dated back to 1899 when Bert Hite, a researcher in an Agriculture Research Station in Morgantown, West Virginia (Tao et al., 2014). Hite’s intention in using this technology was to pasteurize milk as well as other food products (Rivalain et al., 2010). In the year 1914, Hite demonstrated results that fungus (yeasts) and lactic acid microbes are both demanding constituents in sweetened, fully developed fruit were extra sensitive to pressure alternatively to living things, specifically endospore-forming bacilli correlated alongside of vegetables (Rivalain et al., 2010). HPP can be managed in circulating (ambient) or refrigerated temperatures; which allows for the eradication of thermally originated cooked abnormal flavors and tastes (Nath et al., 2019).

A vast number of food research professionals and nutritionists have largely designed and investigated projects in examining the effect of HPP on pastes and puree products (Nath et al., 2019). Some of the investigated products included, peach, pumpkin, nectarine, apple, strawberry, blackberry, tomato, and carrot purees (Nath et al., 2019). Presently, there are no publications that explore toxicity effects of the HPP-treated food commodities; even though reactions to certain food materials were discussed (Hugas et al., 2002; Oey et al., 2008; Nath et al., 2019). There are four major fundamental principles that regulate the outcome of HPP on food elements. The principles include, Le Chatelier’s principle, Isostatic pressing, Microscopic ordering principle, and Arrhenius relationship. The first principle Le Chatelier discussed that ‘any reaction, conformational change, or phase transition which is accompanied by a decrease in volume is enhanced under high-pressure conditions (Yordanov and Angelova, 2010; Arora and Chauhan, 2019) and the processes which involve volume increase are inhibited by pressure (Butz and Tauscher, 1998; Arora and Chauhan, 2019). At low temperatures, such as those ranging between 0 and 40°C the HPP equipment does not have an effect on the covalent bonds. However, water fearing (hydrophobic) and ionic synergism are culpable for the formation of tertiary and quaternary molecular structures; which are normally modified at pressures exceeding 200 MPa (Hendrickx et al., 1998; Arora and Chauhan, 2019).

The second principle, Isostatic pressing commonly referred to as Pascal’s Principle, within the HPP process food products undergo a series of condensed and uncondensed consistently from any direction that is inattentive to dimensions and structures of the chosen food products dissimilar to thermal processing (Yaldagard et al., 2008 ; Huppertz, 2010; Arora and Chauhan, 2019). Even though this principle conserves food products from devastation and deshaped (Arora and Chauhan, 2019). Even though proteins are malleable and compressible, globular proteins undergoing HPP have the potential to become denatured because of empty space on the interior of their molecular structures; which makes them susceptible to compressibility (Arora and Chauhan, 2019).

The third principle, Microscopic ordering principle states that at consistent temperatures, an escalation in the overall pressure will escalate the intensity of molecular ordering of specific substances (Arora and Chauhan, 2019). At elevated temperatures, particles demonstrate reduced disorder which allows chemical responses to occur (Arora and Chauhan, 2019). The fourth principle, Arrhenius relationship, displays numerous response rates that were directly affected by the HPP processing methods (Arora and Chauhan, 2019). The net effects cab be harmonious, preservative, or combative (Balasubramaniam et al., 2015 ; Arora and Chauhan, 2019).

Japan was the first country to test, design, and initiate the HPP technology (Arora and Chauhan, 2019). This technology moved from Japan to American, Europe, and select Asian countries (Arora and Chauhan, 2019). There are a total of ten countries that have production facilities of HPP machines, to include the United States of America, Japan, China, Spain, France, Poland, Germany, Netherlands, United Kingdom, and Sweden (Arora and Chauhan, 2019). These countries produce, sell, and loan these machines to different companies as well to conduct different studies. Within the United States there are a total of four manufacturing facilities; which are Avure Technologies, Engineered Pressure Systems Incorporated, Harwood Engineering, and Elmhurst Research (Arora and Chauhan, 2019).

### Literature Review

Foodborne *Salmonella* infections are a significant public health concern in the United States and also worldwide. There are a number of factors that contribute to food poisoning, however; in many cases it is caused by different bacteria that can affect both high risk and healthy individuals. In reference to high risk individuals, they are considered to be a population termed YOPI; which stands for the Young, Old, Pregnant, and Immunocompromised (Bajpai et al., 2012). Around 30% of the total United States population are considered to at risk for foodborne diseases. An additional concern is the increase in Salmonellosis; which is a zoonotic disease caused by *Salmonella* and primarily transmitted through food products. The Centers for Disease Control and Prevention (CDC) estimates that *Salmonella* causes about 1.2 million illnesses, 23,000 hospitalizations, and 450 death annually in the United States (CDC, 2019a). Some current outbreaks related to *Salmonella* include Tahini products, Raw chicken products, Ground Beef, Raw Turkey products, Pet Hedgehogs, and Pet Guinea Pigs (CDC, 2019b). The most current and severe cases of *Salmonella* are from non-typhoidal; which accounts for 1, 095,079 illness episodes, equivalent to a 27.2% hospitalization rate, and a 0.5% death rate (Cummings et al., 2016). The number of years of life lost (YLL) is 17200; which is second to toxoplasmosis. In terms of DALY, *Salmonella* infections are calculated based on years spent with disability, specific time lived with post-infectious irritable bowel syndrome PI-IBS (Scallan et al., 2015). Scallan et al., 2011 and Hoffmann et al., 2012 both provided QALY background information and comparisons between the top pathogens of concern.

The quality-adjusted life years (QALYs) which equates to one year in perfect health are among the integrated measures of disease burden and it allows comparison of burden from diseases with different health results. These comparisons are critical to risk-based food safety policy. Nontyphoidal *Salmonella enterica* and other pathogens (*Campylobacter spp*., *Clostridium perfringens, Cryptosporidium parvum, Cyclospora cayetanensis, Escherichia coli* O157:H7, non-O157 Shiga toxin–producing *E. coli, Listeria monocytogenes, Shigella, Toxoplasma gondii, Vibrio vulnificus, Vibrio parahaemolyticus* and other noncholera Vibrio, and *Yersinia enterocolitica*) cause over 95% of annual foodborne illnesses, hospitalizations, and deaths attributable to U.S. foodborne disease of specified etiology. More extensively, the estimated QALY loss per death for this pathogen is 43.0 and it is the third highest among other pathogens studied, with *Shigella spp*., and *Yersinia enterocolitica* ranking first and second highest, respectively. In addition, the average QALY loss per death for children less than one year old (infants) is 67.0 while for individuals seventy and above (elderly) is 2.6 with NTS accounting for 13.8% and 6.6%, respectively. The total average QALY loss per death between the age category of < 1 and 70^+^ is 39.8 and NTS having a mean of 9.1% (Batz et al 2014).

Shiga-toxin producing *Escherichia coli* O157:H7 accounts for approximately 176,000 illnesses, 3,700 hospitalizations, and 30 deaths in the United States (Scharff 2015; Scallan et al., 2011). It accounts for over 400 serotypes that affect individuals healthy and those a part of the YOPI population (Scheutz and Stockbine, 2005). There are two major cytotoxins that *Escherichia coli* O157:H7 produces and they are Stx1 and Stx2; which are encoded by stx genes carried via bacteriophages (Cha et al., 2018; O’Brien et al., 1984). These Shiga toxins, also known as verotoxins are associated with infant diarrhea, haemolytic uraemia syndrome (HUS), haemorrhagic colitis, thrombotic thrombocytic purpura (Kovacs et al., 1990; Smith and Scotland., 1988; Tesh and O’Brien 1991). Cattle have been known to be reservoirs of many STEC serotypes that cause illness and infection (Nataro and Kaper, 1998).

Shiga-toxin producing *Escherichia coli* Non-O157 have had a vast number of isolates that have been known to cause illness and outbreak. A total of six O groups; which is comprised of thirteen *serotypes have been identified by the Centers for Disease Control and Prevention (CDC) as a* cause of 71% of diseases (Bosilevac and Koohmaraie, 2011). The exact break down of the 71% as described by *Bosilevac and Koohmaraie, 2011*, was 22% O26, 16% O111, 12% O103, 8% O121, 7% O45, and 5% O145. In the United States annually, the burdens of illness and disease related to non-0157 is doubled in comparison to Shiga-toxin producing *Escherichia coli* (STEC) (Scallan et al., 2011) and the related serogroup strains are O26, O45, O103, O111, O121, and O145 (CDC, 2016).

Shiga-toxin producing *Escherichia coli* Non-O157 according to the CDC surpasses the number of illnesses and complications related to STEC O157 (Hadler et al., 2011; Hale et al., 2012; Valilis et al., 2018). Additionally, it was discussed that STEC Non-O157 is vastly undetected in many cases of outbreak (Hadler et al., 2011; Hale et al., 2012; Valilis et al., 2018). The two most common post infection complications associated with STEC Non-O157 are dysentery and hemolytic uremic syndrome (HUS) (Valilis et al., 2018). The most common symptom associated with non-O157 STEC is diarrhea; which has been observed in non-outbreaks as well as outbreaks of this pathogen (Clogher et al., 2012; Valilis et al., 2018). *Valilis et al*., *2018* determined 129 varying non-O157 O serogroups of STEC directly related with analytical episodes of diarrhea.

*Listeria monocytogenes* is a psychotropic ‘cold-loving’ bacteria that is typically associated with luncheon (deli, cold cuts, sandwhich) meats. Deli meat is typically pre-cooked (pre-processed), wrapped in materials suitable for both warm and dry climate conditions; as it sometimes will endure extended refrigeration and freezer storage periods (Xu et al., 2017). Due to the fact that this bacterium is psychotropic, it can proliferate to menacing levels in suitable refrigeration conditions (Xu et al., 2017; Zhu et al., 2005).

*L. monocytogenes* has the capability to enter a vast number of different mammalian cell types. This is possible because the *L. monocytogenes* pathogen subjects the plasma membrane portion of the cell to reshaping; which allows bacterial engulfment to take place (Pizarro-Cerda et al., 2018). Following the cell invasion, the internal vacuole portion of the cell is subjected to translocation to the cytoplasm portion of the cell (Pizarro-Cerda et al., 2018). Once the translocation to the cytoplasm occurs, bacterial duplication occurs (Pizarro-Cerda et al., 2018). To move between cells *L. monocytogenes* utilizes an actin established maneuverability scheme that grants cytoplasmic bacterial locomotion (Pizarro-Cerda et al., 2018). *L. monocytogenes* infections also effect ion transportation, protein interaction, post-translational state alterations, manufacturing of phosphoinositide, inherent immune feedback, gene appearance, and Deoxyribonucleic acid cohesion (Pizarro-Cerda et al., 2018).

*L. monocytogenes* has the capability to propagate over a wide scope of detrimental physical conditions to include low temperature and pH and salt elevated conditions (Olaimat et al., 2018). Three major post complications are possible if someone contracts *L. monocytogenes*, and they are Septicemia, meningitis, and meningoencephalitis all typically effect individuals with compromised immune systems (Scallan et al., 2011; Swaminathan and Gerner-Smidt, 2007). In newborns intrusive viruses are possible and heighten the potential for stillbirths and natural abortions (Scallan et al., 2011; Swaminathan and Gerner-Smidt, 2007). In terms of treatment options, *L. monocytogenes* has demonstrated resistance to five antibiotics that were previously prescribed by physicians prior to the year 2018 (Olaimat et al., 2018).

### Apple Cider

Apple’s scientific name is *Malus domestica* and it is rich in numerous bioactive compounds; which include, antioxidants, phenolic acids, flavonoids, and flavonols (Roobab et al., 2018). The phenolic composition and antioxidant competency are both beneficial from a health aspect (Zielinski et al., 2019). Also, apple juice is rich is carotenoids, which are bioactive phytochemicals that are very common in fruits and vegetables (Galvis-Sánchez and Vinholes, 2018). Carotenoids are also labile amalgmations vulnerable to isomerization and corrosion-related reactions (oxidation) which are advanced in the presence of acids, heat, and light in numerous storage conditions (Galvis-Sánchez and Vinholes, 2018).

One of the fastest ways of destructing bioactive compounds is via traditional thermal treatment methods which are used in appeasing microbial pathogens (Roobab et al., 2018). In addressing issues regarding the effect of traditional thermal treatments; many research studies (Roobab et al., 2018; Azhuvalappil et al., 2010) have demonstrated the effectiveness of apple juice. In the studies mentioned above, the apple juice has maintained its high quality while also reducing microorganisms.

Apple Cider was used as the vehicle for this experiment due to its historical significance in the foodborne industry especially in studies involving *Escherichia coli* O157. In 1991, there was an apple cider outbreak that lead to alertness regarding acidic foods being used as hosts of *Escherichia coli* O157:H7 (Feng, 2012).

There are a vast number of technologies that are currently emerging; however, two of the most important are high hydrostatic pressure (HHP) and high-pressure processing. The food industry is the main one that these technologies are currently used in present-day food processing. One of the main aims is to conserve taste, freshness, and extend shelf life of food products (Loria, 2017; Allison et al., 2018; Rastogi et al., 2010). This technology functions in exalting severe pressure to pathogens to immobilize and make harmful pathogens extinct (Loria, 2017). Serafina Palandech, a CEO and co-founder of an organic chicken nuggets farm, made the following statement regarding HPP technology, ‘HPP is a wonderful, clean method for extending shelf life on natural and organic products without having to add any additives or preservatives.’ Furthermore, the technologies are being tested to replace, or complement conventional interventions in current food processing methods (Allison et al., 2018; Rastogi et al., 2010). This technology is being used as a mean to prevent foodborne diseases, episodes, and mitigate public health concerns of consumers (Allison et al., 2018; Rastogi et al., 2010). The journey of HPP began in academic settings several decades ago (Balasubramaniam et al., 2016); and due to engineering advances to make them commercially available the desire for this technology has rapidly grown (Balasubramaniam et al., 2016; Rastogi et al., 2010). Presently, over 100 food products globally are brought to the market from using HPP technology; which accounts for about $9 billion annually (Allison et al., 2018; Pressure BioScience Inc., 2017) The National Advisory Committee on Microbiological Criteria for Foods (NACMCF) recommended that the term pasteurization be redefined and for HPP to be considered supplementary non-thermal pasteurization (Allison et al., 2018; Wang et al., 2016).

High pressure processing can be done at both extremely hot and extremely cold environments. The typical range of pressures that HPP utilizes is between 100 – 1000 Mpa and quintessentially temperatures under 60°C (Sheen et al., 2015). There is a three-step mechanism that the HPP machine utilizes and it includes, cell membrane alteration (Moussa et al., 2009), ribosome dissociation (Abe, 2007), and cell structure damage/destruction (Hsu et al., 2014). In terms of cold pasteurization, it is a non-thermal form of preservation; which utilizes hydrostatic pressures ranging between 100 – 1000 MPa (Rendueles et al., 2011). HPP involves liquids which is usually water; which serves as the pressure transmission medium to treat food materials. Many of the pressure levels that are commonly used in commercial applications range between 200-600 MPa; and the pressure used depends on the specific intrinsic and extrinsic properties of the product (Mújica-Paz et al., 2011).

## Materials and Methods

Five strain habituated *Salmonella* serovars (ATCC^®^ numbers 13076, 8387, 6962, 9270, 14028); four strain habituated *Listeria monocytogenes* serovars (ATCC ^®^ numbers 51771, 51779, 13932, and 20011L2625); six strain habituated Shiga Toxin-producing *Escherichia Coli* serovars (ATCC^®^ numbers BAA 2196, 2193, 2192,2219, 2215, and 2440); and *Escherichia Coli* Non-O157 serovars (ATCC ^®^ numbers) will be used for inoculation of apple juice. The bacterial strains are Centers for Disease Control and Prevention (CDC) outbreak strains purchased from American Type Culture Collection (ATCC).

For each strain, a loopful from frozen glycerol stock will be aseptically transferred into 10 ml Tryptic Soy Broth plus 0.6% yeast extract (TSBYE) (Difco, Becton, Dickinson and Company, Sparks, Md.), (BeanTown Chemical), and then incubated 20-24 hours at 37°C. One loopful of the above-mentioned overnight suspension will be streak plated onto the surface of Tryptic Soy Agar plus 0.6% yeast extract (TSAYE) (Difco, Becton, Dickinson and company, Sparks, Md.), and incubated at 37°C for 24 hours. The plates will be stored up to a month at 4°C prior to the experiment. Five days prior to the experiment, each strain will be activated by culturing a single colony from the above-mentioned plates stored at 4°C into 10 ml TSBYE, after incubation at 37°C for 24 hours. A 100 μl aliquot will be sub-cultured into 10 ml TSBYE and incubated at 37°C for 24 hours.

Cells will be harvested using centrifugal force at 5,000 RPM (5424 x g) for 15 minutes. After removal of supernatant, in order to remove sloughed cell components, excreted secondary metabolites, and growth media, the cells will be washed with10 ml Phosphate Buffer Saline (PBS, pH 7.4; 0.2g/L KH_2_PO_4_, 1.5g/L Na_2_HPO_4_.7H_2_O, 8.0 g/L NaCl, and 0.2 g/L KCl), and re-centrifuged using the above-mentioned time and intensity, and resuspended in 10 mL Apple Cider. After removal of supernatant, to improve the external validity of the challenge study, prior to experiment, each strain will be individually habituated in sterile apple juice for 72 h at 4°C to allow acclimatization of the pathogen to low temperature and intrinsic factors of the food (Fouladkhah et al., 2012). For the experiment regarding background microflora, a product without any thermal or non-thermal treatment will be used. The five habituated strains of *Salmonella*, four habituated strains of *Listeria*, six STEC strains, and six habituated strains of *E*.*coli* Non-O157 will then be combined, on the day of the experiment and used as inoculum to conduct the microbiological challenge study.

### HPP treatment

Hydrostatic pressure (Barocycler Hub440, Pressure Bioscience Inc., South Easton, MA) of 15,000 to 55,000 PSI (103 to 380 MPa) will be applied at various time intervals for decontamination of the inoculated pathogens. The pressure transmission fluid that will be used in this study is water, and the pressurization time values reported will exclude the time for pressure increase and the release time. The pressure levels and pressurization times will be set manually. The vessel (PULSE tube) containing bacteria and food vehicle will be subjected to pressure treatment at 50,000, 60,000, 70,000, 80,000 and 87,000 PSI (345, 415, 480, 550 and 600 MPa), with a holding time of 3, 4, and 5 minutes at 4°C and 45°C; these experiments will be performed in two replicates per treatment. The unit is equipped with a water jacket and a circulating water bath surrounding the reaction chamber for precise application of hydrostatic pressure at controlled temperature. The challenge experiment will be conducted in Barocycler reaction PULSE Tubes (2 mL), enabling the precise application of pressure at control temperature. The size of the chamber is 40 ml. Internal pressure, temperature, compression rate will be monitored every 3 seconds using the Barocycler HUB 2.3.11 Software. The temperature will be set using a circulating water bath that surrounds the chamber and is monitored using two k-type thermocouples inserted inside the chamber, secured by thermal paste.

### Microbiological and pH analyses

Pressurized and control cell suspensions in PULSE tubes will be neutralized using D/E neutralizing broth (Difco, Becton Dickinson), and serially diluted with 0.1% Maximum Recovery Diluent (MRD) (Difco, Becton Dickinson), and spread plate on TSAYE to enhance the recovery of injured cells (Difco, Becton Dickinson), for enumeration of total aerobic bacteria and *Salmonella* serovars counts after incubation at 37°C for 48 h. The non-selective medium is formulated to enhance the recovery of injured cells based on a preliminary trial in the laboratory.

The pH of substrates will be measured at the time of inoculation with the use of a digital pH meter (Mettler Toledo, AG, Switzerland).

Experimental design and statistical analyses

The experiment was conducted in two microbiologically independent replicates as blocking factors of a randomized complete block design. The study included three independent repetitions per time/temperature/pressure within each block. The data management (calculation of CFU mL and Log CFU/mL) and graphical presentation of the results were analyzed using SAS GLM procedure using Tukey-and Dunnette-adjusted ANOVA and also Microsoft Excel Office package. The statistical analyses was carried out using general linear model and mixed procedures of SAS 9.4 software (SAS Inst., Cary, N.C.). The analyses was conducted at type 1 error level of 5 % (α = 0.05).

## Results and Discussion

### Sensitivity and Inactivation of Shiga Toxin-Producing Escherichia coli at 4°C and 45°C

As demonstrated in Figures 1 and 2, the samples treated at 4 and 45°C, had similar (p ≥ 0.05) temperature values (mean ± SE) before and after the treatments. Across all treatments at 4°C, the values before treatment were 6.05 ± 0.3°C and were 5.18 ± 0.4°C after the treatments. For samples treated 45°C as well, temperature recordings were similar (p< 0.05) before and after treatments. The temperature values were 44.29 ± 0.1 and were 45.02 ± 0.1, before and after treatments respectively.

**Figure 1.**
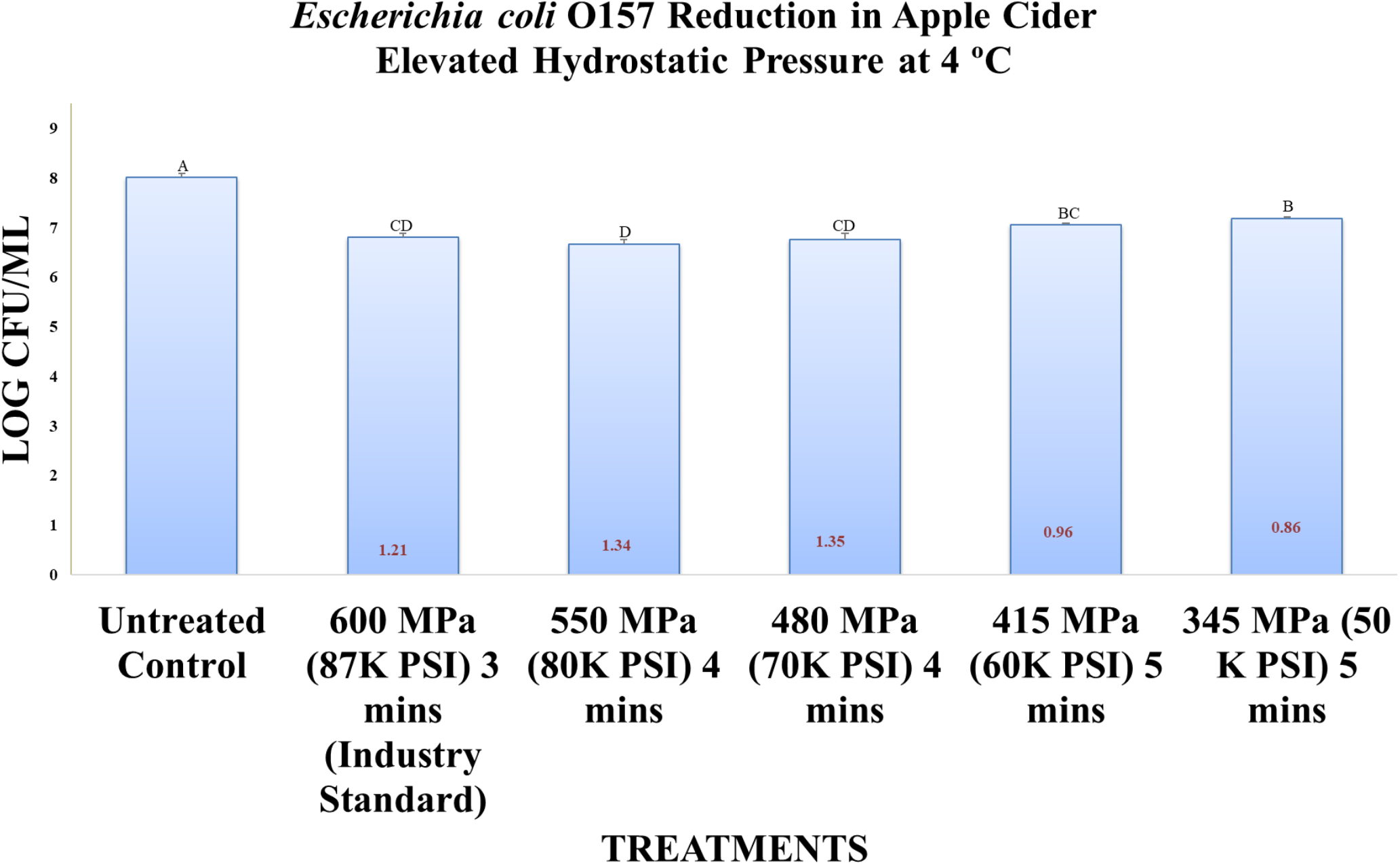
*Escherichia coli* Reduction in Apple Cider Elevated Hydrostatic Pressure at 4°C

**Figure 2.**
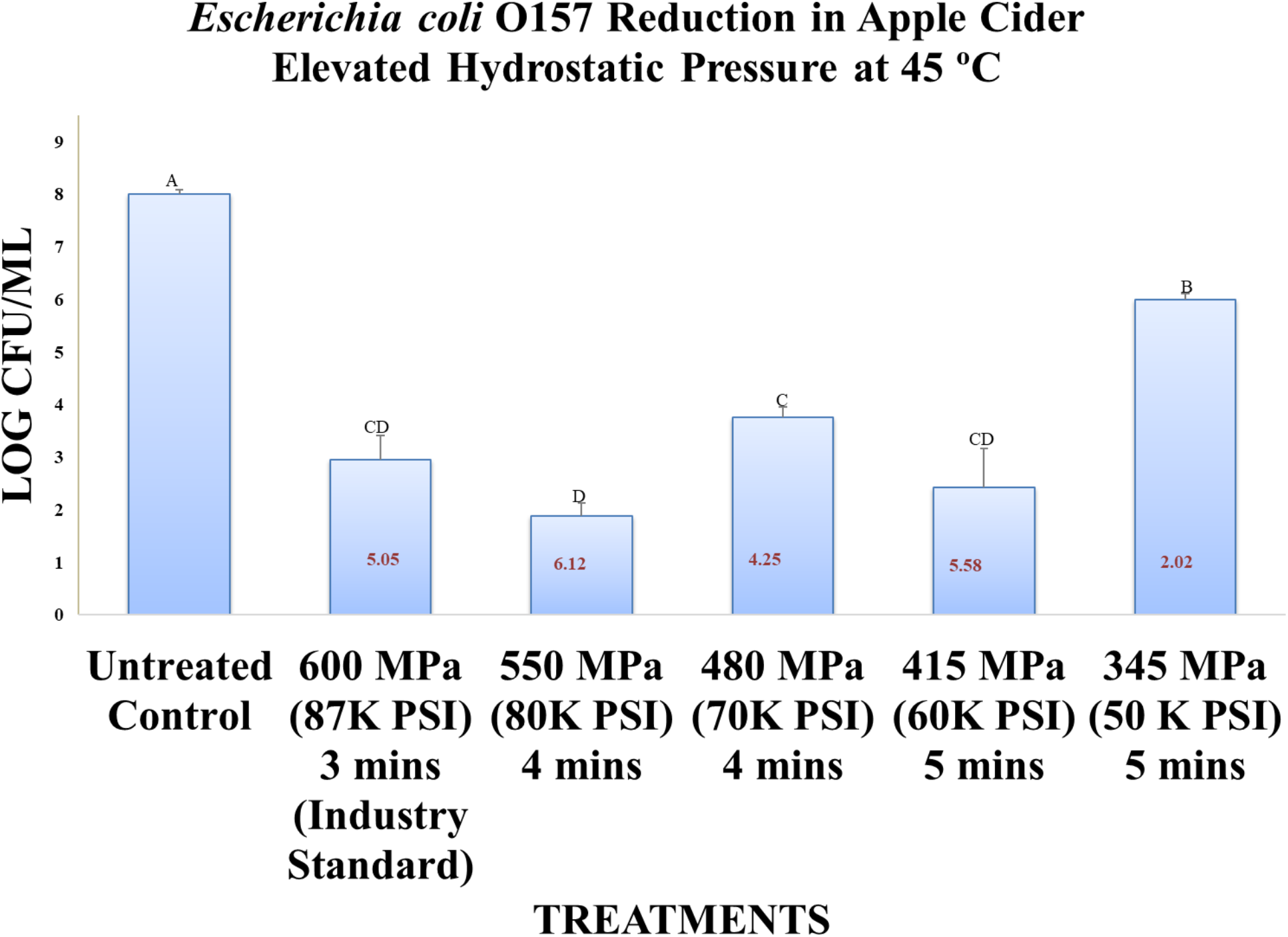
*Escherichia coli* Reduction in Apple Cider Elevated Hydrostatic Pressure at 45°C

The pH levels of the samples were also similar (p ≥ 0.05) before and after treatments. For samples treated at 4°C, and prior to neutralization, the pH value (mean ± SD) and range were from 3.86 ± 0.1and 4.53 ± 0.2, respectively. After neutralization, these values were expectedly increased (p < 0.05) to 6.36 ± 0.1 and 6.85 ± 0.1. Similarly, for samples treated at 45°C, these values were 3.82 ± 0.2 to 4.42 ± 0.2, and 6.38 ± 0.3 to 7.45 ± 0.3, before and after neutralization.

Counts of the pathogen for untreated controls were 8.01 ± 0.1 log CFU/mL (mean ± SD). Treatments for three min at 600 MPa resulted in reduction (p < 0.05), specifically 6.80 ± 0.2. Treated samples at 550 MPa, 480 MPa, 415 MPa, and 345 MPa after three min, four min, and five min, respectively had counts of 6.67 ± 0.2, 6.76 ± 0.3, 7.05 ± 0.1, and 7.18 ± 0.1 log CFU/mL. Longer duration of pressure treatments, predictably resulted in higher inactivation of the pathogen. At lower pressures and higher treatment times for Shiga toxin-producing *Escherichia coli* were reduced (p < 0.05).

The pathogen counts were further reduced (p< 0.05) to <1.21 ± 0.2, <1.34 ± 0.2, <1.25 ± 0.3, <0.96 ± 0.1, and <0.83 ± 0.1 log CFU/mL, for 3-min, 4-min, and 5-min treatments at 600, 550, 480, 415, and 345 MPa and 4°C (mean ± SD), respectively. Even mild pressure treatments, coupled with, elevated heat resulted in an appreciable reduction (p < 0.05) of Shiga toxin-producing *Escherichia coli*. These counts were higher (p >0.05) at 45°C relative to those samples treated at 4°C.

Counts of the pathogen for untreated controls at 45°C were 8.00 ± 0.1 log CFU/mL (mean ± SE). Pathogen counts for the above five pressures, 600 MPa, 550 MPa, 480 MPa, 415 MPa, and 345 MPa at 45°C were 2.96 ± 0.4, 1.89 ± 0.2, 3.76 ± 0.2, 2.43 ± 0.7, and 5.99 ± 0.1 log CFU/mL (mean ± SE), respectively. The pathogen counts were further reduced (p< 0.05) to 5.05 ± 0.4, 6.12 ± 0.2, 4.25 ± 0.2, 5.58 ± 0.7, and 2.02± 0.1log CFU/mL, for 3-min, 4-min, and 5-min treatments at 600, 550, 480, 415, and 345 MPa and 45°C (mean ± SD), respectively.

### Sensitivity and Inactivation of Non-Shiga Toxin-Producing Escherichia coli(nSTEC) at 4°C and 45°C

As demonstrated in Figures 3 and 4, the samples treated at 4 and 45°C, had similar (p ≥ 0.05) temperature values (mean ± SE) before and after the treatments. Across all treatments at 4°C, the values before treatment were 5.51 ± 0.1°C and were 5.53 ± 0.1°C after the treatments. For samples treated 45°C as well, temperature recordings were similar (p< 0.05) before and after treatments. The temperature values were 44.90 ± 0.2 and were 45.75 ± 0.2, before and after treatments respectively.

**Figure 3.**
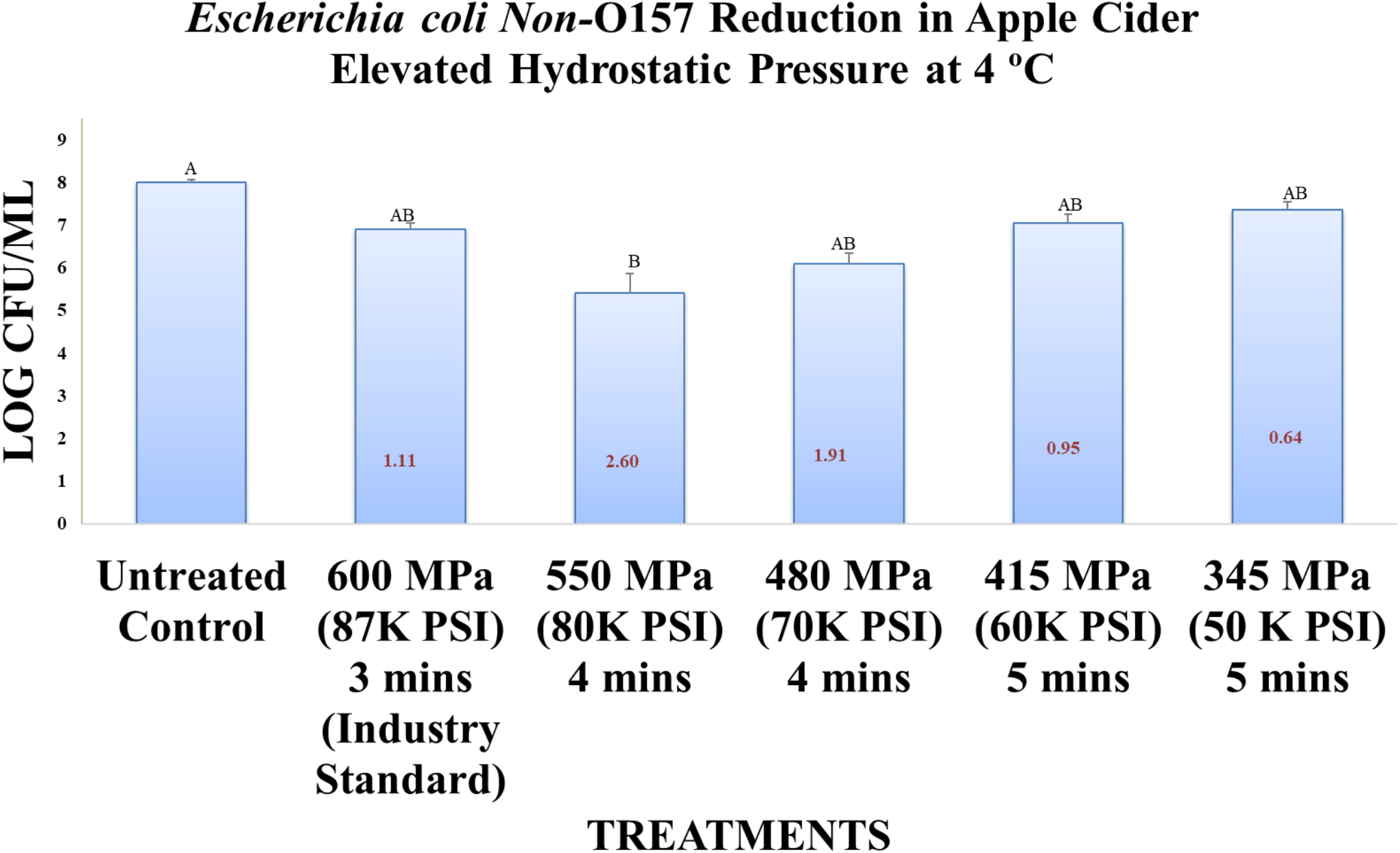
*Escherichia coli* Non-O157 Reduction in Apple Cider Elevated Hydrostatic Pressure at 4°C

**Figure 4.**
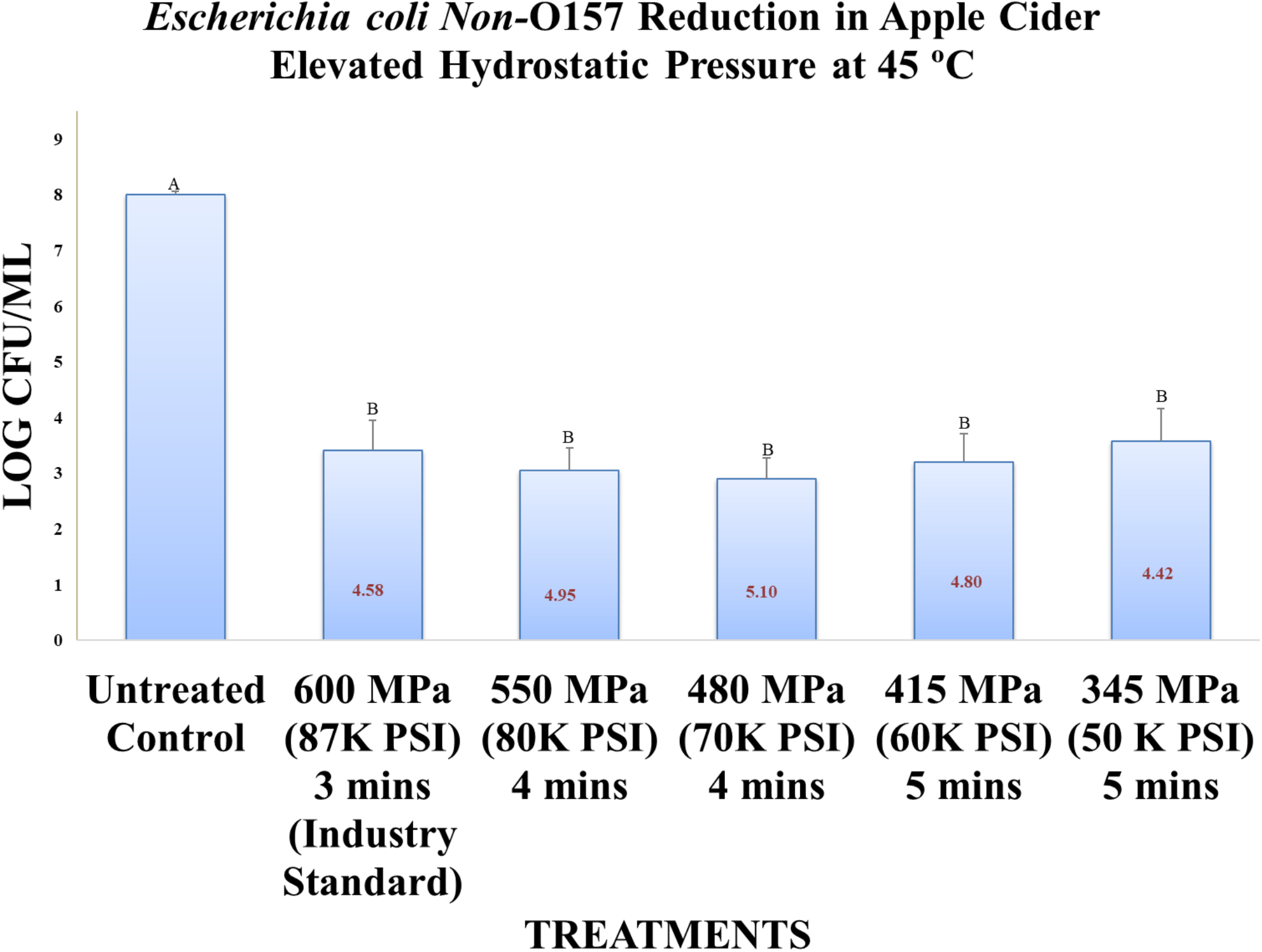
*Escherichia coli* Non-O157 Reduction in Apple Cider Elevated Hydrostatic Pressure at 45°C

The pH levels of the samples were also similar (p ≥ 0.05) before and after treatments. For samples treated at 4°C, and prior to neutralization, the pH value (mean ± SD) and range were from 3.84 ± 0.1and 4.20 ± 0.1, respectively. After neutralization, these values were expectedly increased (p < 0.05) to 6.94 ± 0.1 and 7.39 ± 0.1. Similarly, for samples treated at 45°C, these values were 3.75 ± 0.1 to 4.17 ± 0.1, and 6.74 ± 0.1 to 7.04 ± 0.1, before and after neutralization.

Counts of the pathogen for untreated controls were 8.00 ± 0.1 log CFU/mL (mean ± SD). Treatments for three min at 600 MPa resulted in reduction (p < 0.05), specifically 6.88 ± 0.4. Treated samples at 550 MPa, 480 MPa, 415 MPa, and 345 MPa after three min, four min, and five min, respectively had counts of 5.40 ± 1.1, 6.09 ± 0.6, 7.05 ± 0.5, and 7.40 ± 0.5 log CFU/mL. Longer duration of pressure treatments, predictably resulted in higher inactivation of the pathogen. At lower pressures and higher treatment times for non-Shiga toxin-producing *Escherichia coli* were reduced (p < 0.05).

The pathogen counts were further reduced (p< 0.05) to <1.12 ± 0.4, <2.60 ± 1.1, <1.91 ± 0.6, <0.95 ± 0.3, and <0.60 ± 0.5 log CFU/mL, for 3-min, 4-min, and 5-min treatments at 600, 550, 480, 415, and 345 MPa and 4°C (mean ± SD), respectively. Even mild pressure treatments, coupled with, elevated heat resulted in an appreciable reduction (p < 0.05) of non-Shiga toxin-producing *Escherichia coli*. These counts were higher (p >0.05) at 45°C relative to those samples treated at 4°C.

Counts of the pathogen for untreated controls at 45°C were 8.00 ± 0.1 log CFU/mL (mean ± SE). Pathogen counts for the above five pressures, 600 MPa, 550 MPa, 480 MPa, 415 MPa, and 345 MPa at 45°C were 3.42 ± 0.5, 3.05 ± 0.4, 2.90 ± 0.4, 3.20 ± 0.5, and 3.60 ± 0.6 log CFU/mL (mean ± SE), respectively. The pathogen counts were further reduced (p< 0.05) to <4.58± 0.5, <4.95 ± 0.4, <5.10 ± 0.4, <4.8 ± 0.5, and <4.40 ± 0.6 log CFU/mL, for 3-min, 4-min, and 5-min treatments at 600, 550, 480, 415, and 345 MPa and 45°C (mean ± SD), respectively.

### Sensitivity and Inactivation of Salmonella serovars at 4°C and 45°C

As demonstrated in Figures 5 and 6, the samples treated at 4 and 45°C, had similar (p ≥ 0.05) temperature values (mean ± SE) before and after the treatments. Across all treatments at 4°C, the values before treatment were 6.02 ± 0.3°C and were 4.88 ± 0.4°C after the treatments. For samples treated 45°C as well, temperature recordings were similar (p< 0.05) before and after treatments. The temperature values were 44.66 ± 0.2 and were 45.33 ± 0.1, before and after treatments respectively.

**Figure 5.**
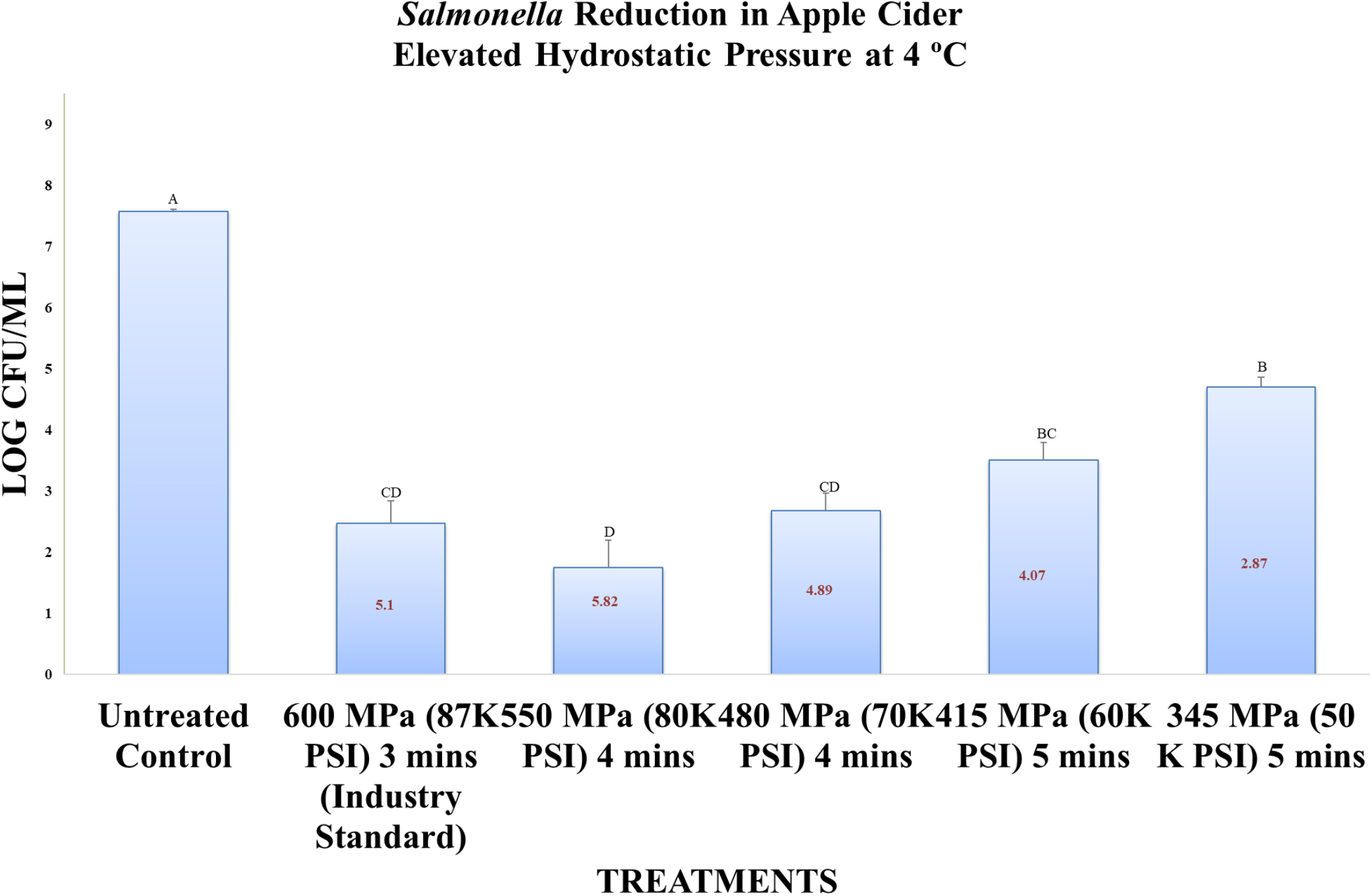
*Salmonella* Reduction in Apple Cider Elevated Hydrostatic Pressure at 4°C

**Figure 6.**
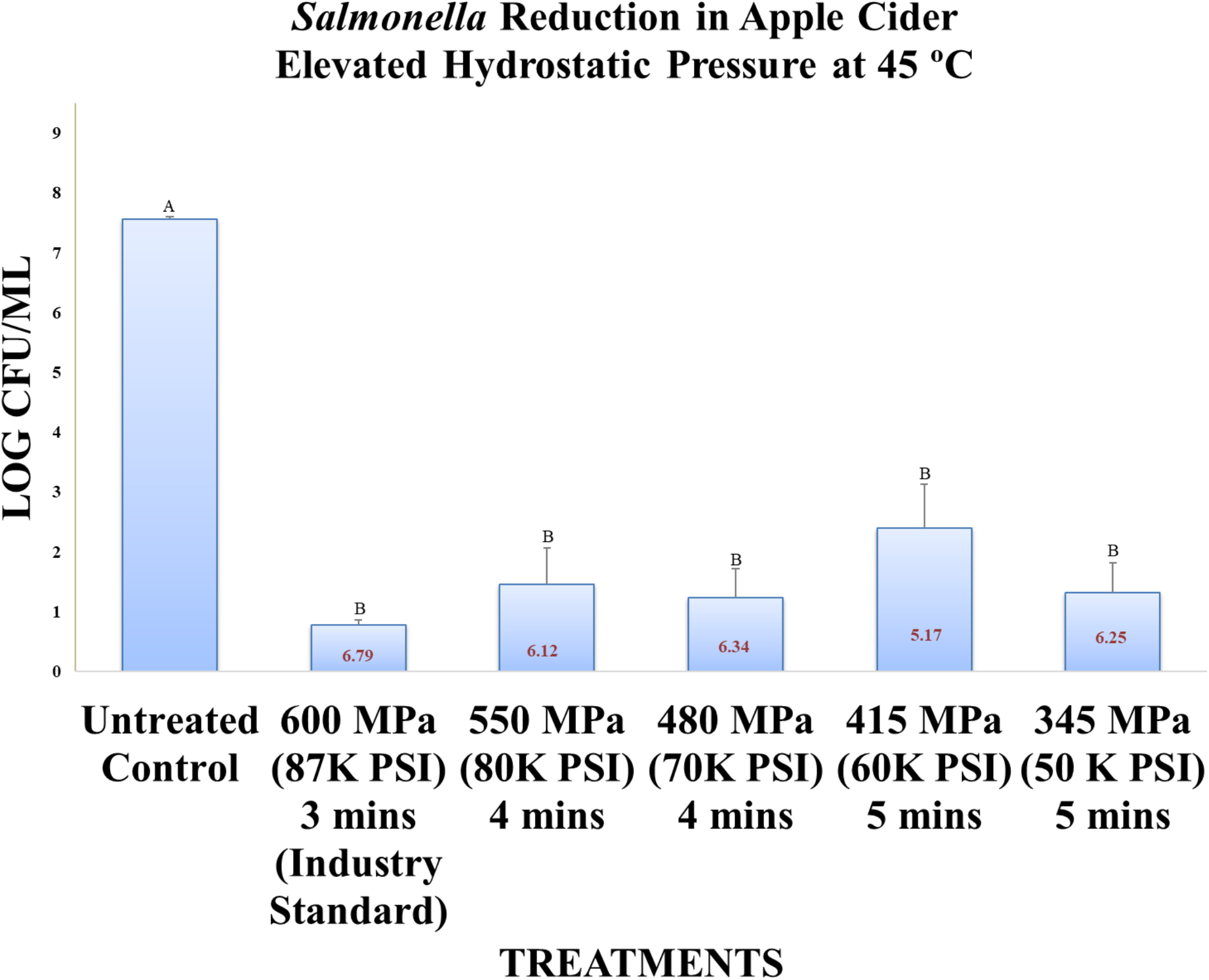
*Salmonella* Reduction in Apple Cider Elevated Hydrostatic Pressure at 45°C

The pH levels of the samples were also similar (p ≥ 0.05) before and after treatments. For samples treated at 4°C, and prior to neutralization, the pH value (mean ± SD) and range were from 3.84 ± 0.2 and 4.85 ± 0.3, respectively. After neutralization, these values were expectedly increased (p < 0.05) to 6.39 ± 0.1 and 7.66 ± 0.3. Similarly, for samples treated at 45°C, these values were 3.80 ± 0.2 to 4.30 ± 0.2, and 6.39 ± 0.4 to 7.51 ± 0.4, before and after neutralization.

Counts of the pathogen for untreated controls were 7.57 ± 0.1 log CFU/mL (mean ± SD). Treatments for three min at 600 MPa resulted in reduction (p < 0.05), specifically 2.47 ± 0.9. Treated samples at 550 MPa, 480 MPa, 415 MPa, and 345 MPa after three min, four min, and five min, respectively had counts of 1.75 ± 1.1, 2.68 ± 0.7, 3.50 ± 0.7, and 4.70 ± 0.4 log CFU/mL. Longer duration of pressure treatments, predictably resulted in higher inactivation of the pathogen. At lower pressures and higher treatment times for *Salmonella* serovars were reduced (p < 0.05).

The pathogen counts were further reduced (p< 0.05) to <5.10± 0.9, <5.82 ± 1.1, <4.89 ± 0.7, <4.07 ± 0.7, and <2.87 ± 0.7 log CFU/mL, for 3-min, 4-min, and 5-min treatments at 600, 550, 480, 415, and 345 MPa and 4°C (mean ± SD), respectively. Even mild pressure treatments, coupled with, elevated heat resulted in an appreciable reduction (p < 0.05) of *Salmonella* serovars. These counts were higher (p >0.05) at 45°C relative to those samples treated at 4°C.

Counts of the pathogen for untreated controls at 45°C were 7.57 ± 0.0 log CFU/mL (mean ± SE). Pathogen counts for the above five pressures, 600 MPa, 550 MPa, 480 MPa, 415 MPa, and 345 MPa at 45°C were 0.78 ± 0.1, 1.45 ± 0.6, 1.23 ± 0.5, 2.40 ± 0.7, and 1.32 ± 0.5 log CFU/mL (mean ± SE), respectively. The pathogen counts were further reduced (p< 0.05) to <6.79± 0.1, <6.12 ± 0.6, <6.34 ± 0.5, <5.17 ± 0.7, and <6.25 ± 0.5 log CFU/mL, for 3-min, 4-min, and 5-min treatments at 600, 550, 480, 415, and 345 MPa and 45°C (mean ± SD), respectively.

### Sensitivity and Inactivation of Listeria monocytogenes at 4°C and 45°C

As demonstrated in Figures 7 and 8, the samples treated at 4 and 45°C, had similar (p ≥ 0.05) temperature values (mean ± SE) before and after the treatments. Across all treatments at 4°C, the values before treatment were 5.43 ± 0.1°C and were 5.47 ± 0.1°C after the treatments. For samples treated 45°C as well, temperature recordings were similar (p< 0.05) before and after treatments. The temperature values were 45.18 ± 0.2 and were 45.76 ± 0.2, before and after treatments respectively.

**Figure 7.**
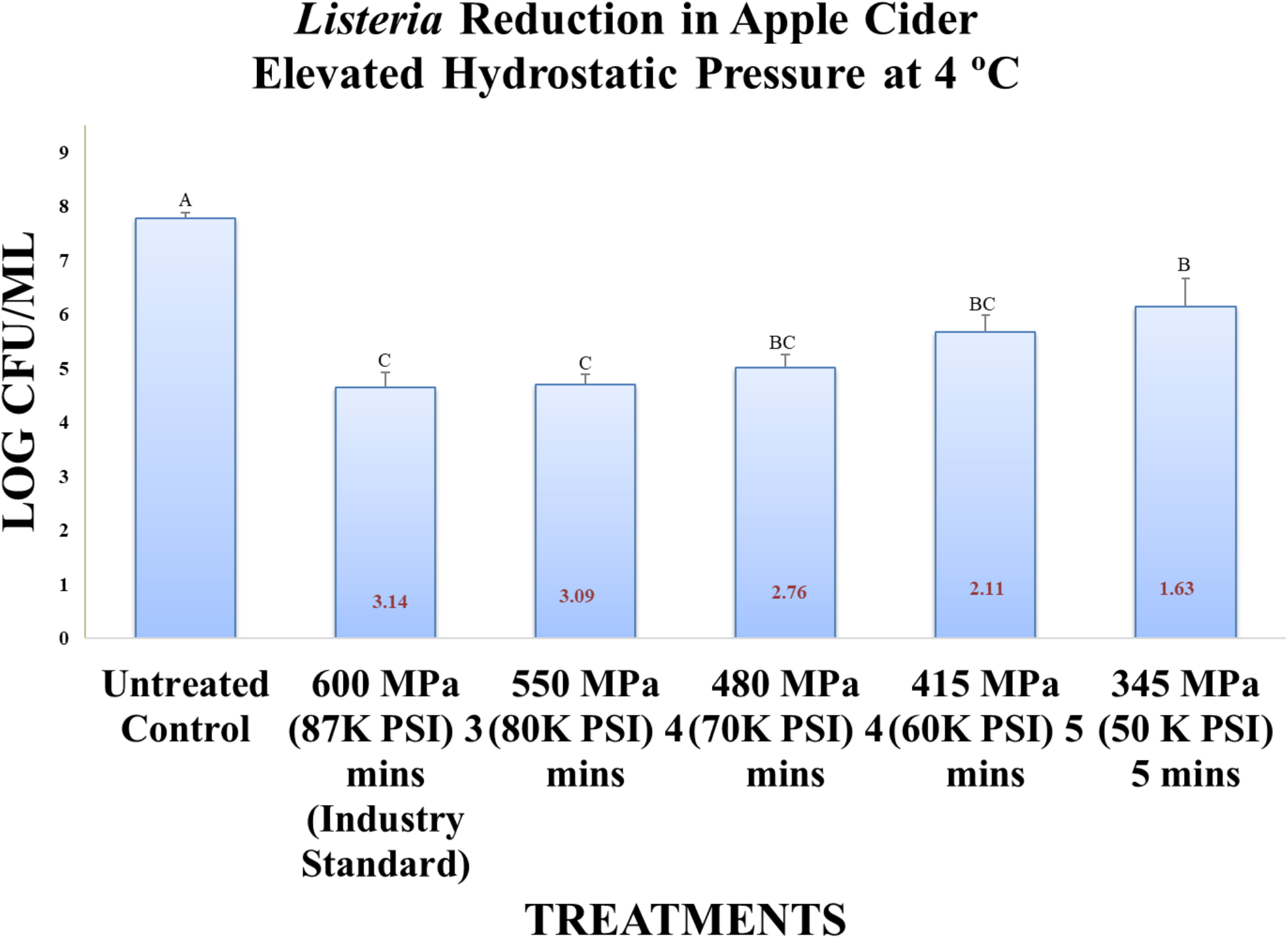
*Listeria monocytogenes* Reduction in Apple Cider Elevated Hydrostatic Pressure at 4°C

**Figure 8.**
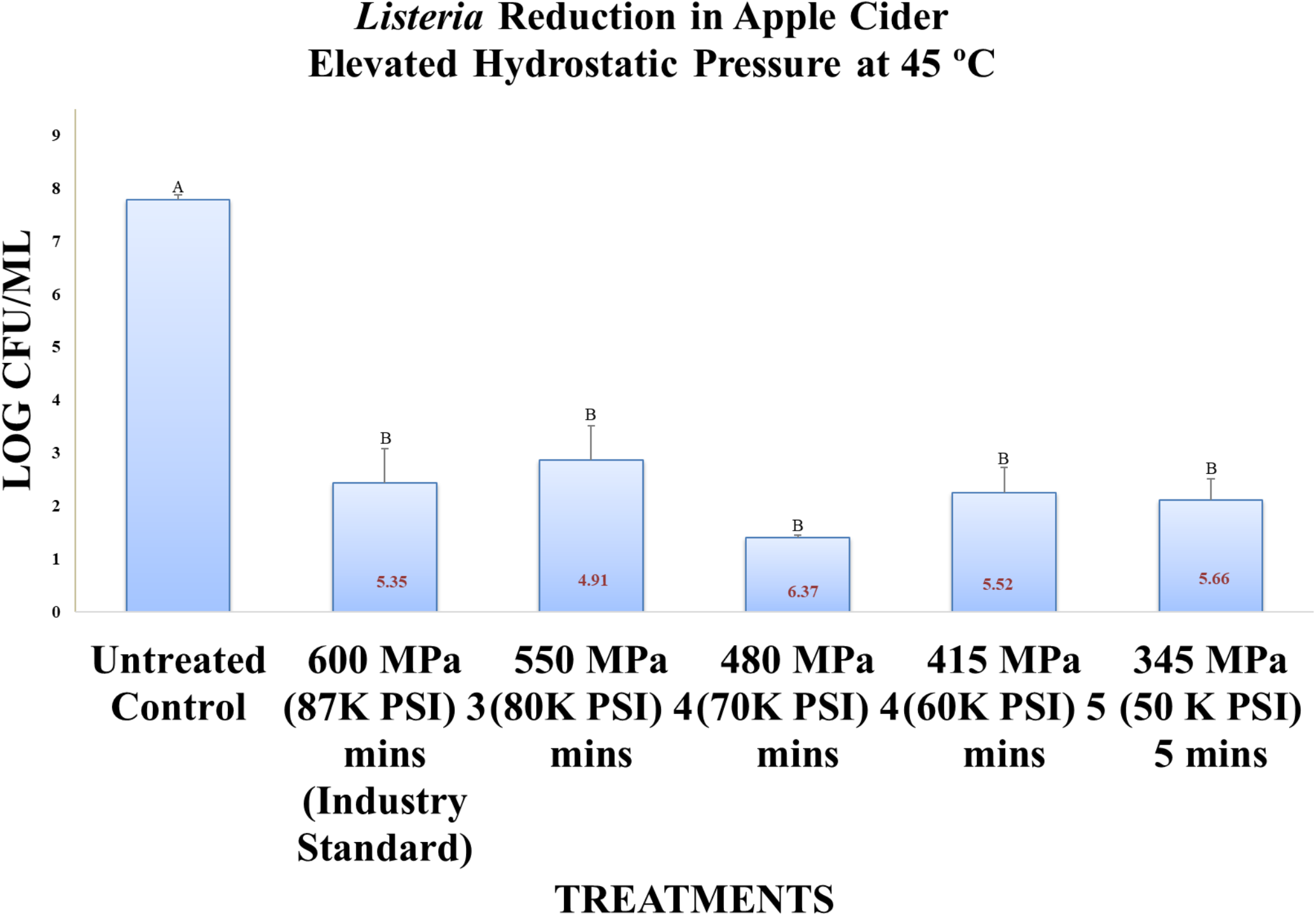
*Listeria monocytogenes* Reduction in Apple Cider Elevated Hydrostatic Pressure at 45°C

The pH levels of the samples were also similar (p ≥ 0.05) before and after treatments. For samples treated at 4°C, and prior to neutralization, the pH value (mean ± SD) and range were from 3.83 ± 0.1 and 4.03 ± 0.1, respectively. After neutralization, these values were expectedly increased (p < 0.05) to 6.94 ± 0.0 and 7.15 ± 0.1. Similarly, for samples treated at 45°C, these values were 3.79 ± 0.1 to 4.02 ± 0.1, and 6.71 ± 0.1 to 7.06 ± 0.0, before and after neutralization.

Counts of the pathogen for untreated controls were 8.00 ± 0.1 log CFU/mL (mean ± SD). Treatments for three min at 600 MPa resulted in reduction (p < 0.05), specifically 4.64 ± 0.7. Treated samples at 550 MPa, 480 MPa, 415 MPa, and 345 MPa after three min, four min, and five min, respectively had counts of 4.70 ± 0.5, 5.02 ± 0.6, 5.67 ± 0.8, and 6.14 ± 1.3 log CFU/mL. Longer duration of pressure treatments, predictably resulted in higher inactivation of the pathogen. At lower pressures and higher treatment times for *Listeria monocytogenes* were reduced (p < 0.05).

The pathogen counts were further reduced (p< 0.05) to <3.36 ± 0.3, <3.30 ± 0.5, <2.98 ± 0.6, <2.33 ± 0.8, and <1.86 ± 1.3 log CFU/mL, for 3-min, 4-min, and 5-min treatments at 600, 550, 480, 415, and 345 MPa and 4°C (mean ± SD), respectively. Even mild pressure treatments, coupled with, elevated heat resulted in an appreciable reduction (p < 0.05) of *Listeria monocytogenes*. These counts were higher (p >0.05) at 45°C relative to those samples treated at 4°C.

Counts of the pathogen for untreated controls at 45°C were 8.00 ± 0.1 log CFU/mL (mean ± SE). Pathogen counts for the above five pressures, 600 MPa, 550 MPa, 480 MPa, 415 MPa, and 345 MPa at 45°C were 2.43 ± 0.7, 2.90 ± 0.6, 1.41 ± 0.0, 2.25 ± 0.5, and 2.11 ± 0.4 log CFU/mL (mean ± SE), respectively. The pathogen counts were further reduced (p< 0.05) to <5.57± 0.7, <5.10 ± 0.6, <6.59 ± 0.0, <5.75 ± 0.5, and <5.89 ± 0.4 log CFU/mL, for 3-min, 4-min, and 5-min treatments at 600, 550, 480, 415, and 345 MPa and 45°C (mean ± SD), respectively.

## Acknowledgments

This study was supported in part by United States Department of Agriculture and Pressure BioScience Inc. Contributions of Sadiya Arias and Kaleh Karim, students of Public Health Microbiology Laboratory of Tennessee State University and technical support of James Behnke and Dr. Nate Lawrence from Pressure BioScience Inc. are sincerely appreciated by the corresponding author of this study. We would also appreciate the feedback of anonymous peers who reviewed this study. We would also like to extend appreciation to Dr. Aliyar Fouladkhah for his assistance on this project too.

## Author Contributions

Anita Scales: Graduate Research Assistant, Tennessee State University. Conducted the main and preliminary laboratory experiments, assisted in data management and analyses, co-wrote the first version of manuscript in partial fulfillment of her doctoral dissertation. Abimbola Allison: Professor, Tennessee State University. Assisted in preparation prior to and when conducting the experiments, and assistance in the preparation of the manuscript. Jayashan Adhikari, Graduate Research Assistant: Assisted during conducting the experiments and also assistance in the preparation of the manuscript. Wendelyn Inman: Associate Professor and Interim Director, Tennessee State University’s Department of Public Health, Health Administration, & Health Sciences program. Assisted in data analysis and preparation of the final manuscript.

## References

1. Abe, F. (2007). Exploration of the effects of high hydrostatic pressure on microbial growth, physiology and survival: perspectives from piezophysiology. Bioscience, Biotechnology, and Biochemistry, 71: 700151–7001511.

2. Allison, A., S. Chowdhury, and A. Fouladkhah. (2018). Synergism of Mild Heat and High-Pressure Pasteurization against Listeria monocytogenes and Natural Microflora in Phosphate-Buffered Saline and Raw Milk. Microorganisms (6), 1–12.

3. Arora, S.K., and O.P. Chauhan. (2019). High-Pressure Processing. Non-thermal Processing of Foods. CRC Press, FL.

4. Azhuvalappil, Z., X. Fan, D.J. Geveke, and H.Q. Zhang. (2010). Thermal and Nonthermal Processing of Apple Cider: Storage Quality Under Equivalent Process Conditions. Journal of Food Quality, 33(5): 612–631.

5. Bajpai, V.K., K. Kwang-Hyun, and S. Chul. (2012). Control of Salmonella in foods by using essential oils: A review. Food Research International, 45(2): 722–734.

6. Balasubramaniam, V.M., S.I.M. Monteagudo, and R. Gupta. (2015). Principles and Application of high-pressure based technologies in the food industry. Annual Review of Food Science and Technology, 6: 435–462.

7. Batz, M., S. Hoffmann, and J. G. Morris Jr. (2014). Disease-outcome trees, EQ-5D scores, and estimated annual losses of quality-adjusted life years (QALYs) for 14 foodborne pathogens in the United States. Foodborne pathogens and Disease, 11(5): 395–402.

8. Bosilevac, J.M., and M. Koohmaraie. (2011). Prevalence and characterization of non-O157 Shiga toxin-producing Escherichia coli isolates from commercial ground beef in the United States. Applied and Environmental Microbiology, 77(6), 2103–2112.

9. Butz, P., and B. Tauscher. (1998). Food Chemistry Under High Hydrostatic Pressure. High Pressure Food Science, Bioscience and Chemistry. The Royal Society of Chemistry, UK, pp. 133–144.

10. Centers for Disease Control and Prevention. (2016). National Enteric Disease Surveillance: Shiga Toxin-producing Escherichia coli (STEC) Annual Report, 2016.

11. Centers for Disease Control and Prevention. (2019a). Salmonella Questions and Answers.

12. Centers for Disease Control and Prevention. (2019b). Reports of Active Salmonella Outbreak Investigations.

13. Cha, W., P.M. Fratamico, L.E. Ruth, A.S. Bowman, J.M. Nolting, S.D. Manning, and J.A. Funk. (2018). Prevalence and characteristics of Shiga toxin-producing Escherichia coli in finishing pigs: implications on public health. International Journal of Food Microbiology, 264: 8–15.

14. Clogher, P., S. Hurd, D. Hoefer, J.L. Hadler, L. Pasutti, S. Cosgrove, S. Segler, M. Tobin-D’Angelo, C. Nicholson, H. Booth, k, Garman, R.K. Mody, and L.H. Gould. (2012). Assessment of physician knowledge and practices concerning Shiga toxin-producing Escherichia coli infection and enteric illness, 2009, Foodborne Diseases Active Surveillance Network (FoodNet). Clinical Infectious Diseases, 54:446–452.

15. Clogher, P., S. Hurd, D. Hoefer, J.L. Hadler, L. Pasutti, S. Cosgrove, S. Segler, M. Tobin-D’Angelo, C. Nicholson, H.B.K. Garman, R.K. Mody, and H. Gould. (2012). Assessment of Physician Knowledge and Practices Concerning Shiga Toxin-Producing Escherichia coli Infection and Enteric Illness, 2009, Foodborne Diseases Active Surveillance Network (FoodNet). Clinical Infectious Diseases, 54 (5):446–452.

16. Feng, P. (2012). Learning from outbreaks of Escherichia coli O157:H7 caused by low pH foods. Case Studies in Food Safety and Authenticity Lessons from Real-Life Situations, A volume in Woodhead Publishing Series in Food Science, Technology, and Nutrition, 244–253.

17. Fouladkhah, A., I. Geornaras, H. Yang, K. Belk, K. Nightingale, D. Woerner, G. Smith, and J. Sofos. (2012). Sensitivity of Shiga Toxin-Producing Escherichia coli, Multi-drug Resistant Salmonella, and Antibiotic-Susceptible Salmonella to Lactic Acid on Inoculated Beef Trimmings. Journal of Food Protection (10), 75–85.

18. Galvis-Sánchez, A.C., and J. Vinholes (2018). Fruit Juices (Apple, Peach, and Pear) and Changes in the Carotenoid Profile. Fruit Juices Extraction, Composition, Quality and Analysis, 5, 59–74.

19. Hadler, J.L., P. Clogher, S. Hurd, Q. Phan, M. Mandour, K. Bemis, and R. Marcus. (2011). Ten-year trends and risk factors for non-O157 Shiga toxin-producing Escherichia coli found through Shiga toxin testing, Connecticut, 2000-2009. Clinical Infectious Diseases, 53(3):269–276.

20. Hale C.R., E. Scallan, A.B. Cronquist, J. Dunn, K. Smith, T. Robinson, S. Lanthrop, M. Tobin-D’Angelo, and P. Clogher. (2012). Estimates of enteric illness attributable to contact with animals and their environments in the United States. Clinical Infectious Diseases, 54(5):472–479.

21. Hendrickx, M., L. Ludikhuyze, I. Van den Broeck, and C. Weemaes. (1998). Effects of high pressure on enzymes related to food quality. Trends in Food Science &Technology, 9:197–203.

22. Hoffmann, S., M.B. Batz, and J.G. Jr. Morris. (2012). Annual cost of illness and quality-adjusted life year losses in the United States due to 14 foodborne pathogens. Journal of Food Protection, 75:1292–1302.

23. Hsu, P. D., E.S. Lander, and F. Zhang. (2014). Development and applications of CRISPR-Cas9 for genome engineering. Cell, 157(6): 1262–1278.

24. Hugas, M., M. Garriga, and J.M. Monfort. (2002). New mild technologies in meat processing: High pressure as a model technology. Meat Science, 62(3): 359–371.

25. Huppertz, T. (2010). High pressure processing of milk. Improving the Safety and Quality of Milk Milk Production and Processing, 1: 373–399.

26. Kovacs, M. J., J. Roddy, S. Gregoire, W. Cameron, L. Eidus, and J. Drouin. (1990). Thrombotic thrombocytic purpura following hemorrhagic colitis due to Escherichia coli 0157:H7. The American Journal of Medicine, 88:177–179.

27. Loria, K. (2017). Under pressure: HPP preserves food, texture and taste. Available at: https://www.fooddive.com/news/food-beverage-high-pressure-processing-hpp/433439/.

28. Moussa, M., V. Espinasse, J. M. Perrier-Cornet, and P. Gervais. (2009). Pressure treatment of Saccharomyces cerevisiae in low-moisture environments. Applied microbiology and biotechnology, 85(1): 165–174.

29. Mújica Paz. H., A. Valdez-Fragoso, C.T. Samson, J. Welti-Chanes, and J.A. Torres. (2011). High-Pressure Processing Technologies for the Pasteurization and Sterilization of Foods. Food and Biopress Technology, 4(6): 969–985.

30. Nataro, J.P., and J.B. Kaper. (1998). Diarrheagenic Escherichia coli. Clinical Microbiology Reviews, 11(1):142–201.

31. Nath, P., S.J. Kale, and B. Bhushan. (2019). Consumer Acceptance and Future Trends of Non-Thermal-Processed Foods. Non-thermal Processing of Foods. CRC Press, FL, pp.

32. O’Brien, A.D., J.W. Newland, S.F. Miller, R.K. Holmes, H.W. Smith, and S.B. Formal. (1984). Shiga-like toxin-converting phages from Escherichia coli strains that cause hemorrhagic colitis or infantile diarrhea. Science, 226 (4675): 694–696.

33. Oey, I., I. Plancken, A. Loey, and M. Hendrickx. (2008). Does high-pressure processing influence nutritional aspects of plant based food system? Trends in Food Science & Technology,19: 300–308.

34. Ohara, E., M. Kawamura, M. Ogino, E. Hoshino, A. Kobayashi, J. Hoshino, A. Yamazaki, and T. Nishiumi (Eds.) K. Akasaka and H. Matsuki. (2015). Application of high-pressure treatment to enhancement of functional components in agricultural products and development of sterilized foods. Subcellular Biochemistry, 72:567–589.

35. Olaimat, A.N., M.A. Al-Holy, H.M. Shahbaz, A.A. Al-Nabulsi, M.H. Abu Ghoush, T.M. Osaili, M.M. Ayyash, and R.A. Holley. (2018). Emergence of Antibiotic Resistance in Listeria monocytogenes Isolated from Food Products: A Comprehensive Review. Comprehensive Reviews in Food Science and Food Safety, 17(5).

36. Olaimat, A.N., M.A. Al-Holy, H.M. Shahbaz, A.A. Al-Nabulsi, M.H.A. Ghoush, T.M. Osaili, M.M. Ayyash, and R. A. Holley. (2018). Emergence of Antibiotic Resistance in Listeria monocytogenes Isolated from Food Products: A Comprehensive Review. Comprehensive Reviews in Food Science and Food Safety, 17(5):1277–1292.

37. Pizarro-Cerda, J., and P. Cossart. (2018). Listeria monocytogenes: cell biology of invasion and intracellular growth. Microbiology Spectrum, 6(6) GPP3-0013-2018.

38. Pizarro-Cerda, J., P. Cossart, (Eds.) V.A. Fischetti, R.P. Novick, J.J. Ferretti, D.A. Portnoy, M. Braunstein, and J.I. Rood. (2018). Listeria monocytogenes: cell biology of invasion and intracellular growth. Microbiology Spectrum, 6(6):3–13.

39. Pressure BioScience Inc. (PBI). (2017). High Pressure Processing. Available online: https://d1io3yog0oux5.cloudfront.net/_57b4ee00ca1d2b0fef872a3c158b2b76/pressurebiosciences/db/399/3062/pdf/Investor+Presentation+Oct+2017.pdf.

40. Rastogi, N.K., L.T. Nguyen, B. Jiang, and V.M. Balasubramaniam. (2010). Improvement in texture of pressure-assisted thermally processed carrots by combined pretreatment using response surface methodology. Food and Bioprocess Technology, 3: 762–771.

41. Rendueles, O., L. Travier, P. Latour-Lambert, T. Fontaine, J. Magnus, E. Denamur, and J.M. Ghigo. Screening of Escherichia coli species Biodiversity Reveals New Biofilm-Associated Antiadhesion Polysaccharides. American Society for Microbiology, mBio, 2: e00043–00011.

42. Rivalain, N., J. Roquain, and G. Demazeau. (2010). Development of high hydrostatic pressure in biosciences: pressure effect on biological structures and potential applications in biotechnologies. Biotechnology Advances, 28:659–672.

43. Roobab, U., R.M. Aadil, G.M. Madni, and A. El-Din Bekhit. (2018). The Impact of Nonthermal Technologies on the Microbiological Quality of Juices: A Review. Comprehensive Reviews in Food Science and Food Safety, 17(2): 437–457.

44. Cummings, P.L., T. Kuo, M. Javanbakht, S. Shafir, M. Wang, and F. Sorvillo. (2016). Salmonellosis Hospitalizations in the United States: Associated Chronic Conditions, Costs, and Hospital Outcomes, 2011, Trends 2000-2011. Foodborne Pathogens and Disease, 13(1):40–48.

45. Scallan, E., M. Hoekstra, B.E. Mahon, T.F. Jones, and P.M. Griffin. (2015). An assessment of the human health impact of seven leading foodborne pathogens in the United States using disability adjusted life years. Epidemiology & Infection, 143:1–10.

46. Scallan, E., R.M. Hoekstra, F.J. Angulo, R.V. Tauxe, M.A. Widdowson, S.L. Roy, J.L. Jones, and P.M. Griffin. (2011). Foodborne Illness Acquired in the United States - Major Pathogens. Emerging Infectious Diseases,17:7–15.

47. Scallan, E., R.M. Hoekstra, F.J. Angulo, R.V. Tauxe, M.A. Widdowson, S.L. Roy, J.L. Jones, and P.M. Griffin. (2011). Foodborne illness acquired in the United States--major pathogens. Emerging Infectious Diseases, 17:7–15.

48. Scharff, R. L., (2015). State estimates for the annual cost of foodborne illness. Journal of Food Protection,78:1064–1071.

49. Scheutz, F., N.A. Strockbine, (Eds.) Brenner, D.J., G.M. Garrity, D.J. Brenner, N.R. Krieg., and J.T. Staley. (2005). Genus I. Escherichia. Bergey’s Manual of Systematic Bacteriology. Springer, NY.

50. Sheen, S., J. Cassidy, B. Scullen, and C. Sommers. (2015). Inactivation of a diverse set of shiga toxin-producing Escherichia coli in ground beef by high pressure processing. Food Microbiology, 52: 84–87.

51. Smith, H.R., and S.M. Scotland. (1988). Vero cytotoxin-producing strains of Escherichia coli. Journal of Medical Microbiology, 26(2):77–85.

52. Swaminathan, B., and P. Gerner-Smidt. (2007). The epidemiology of human listeriosis. Microbes and infection, 9(10):1236–1243.

53. Swaminathan, P. and P. Gerner-Smidt. (2007). The epidemiology of human listeriosis. Microbes and Infection, 9:1236–1243.

54. Tao, Y., D-W. Sun, E. Hogan, and A. Kelly. (2014). High Pressure Processing of Foods: An Overview. Emerging Technologies for Food Processing, 2(1). Academic Press, UK, pp. 3–24.

55. Tesh, V.L., and A. D. O’Brien. (1991). The pathogenic mechanisms of Shiga toxin and the Shiga-like toxins. Molecular Microbiology, 5:1817–1822.

56. Valilis, E., A. Ramsey, S. Sidig, and H.L. DuPont. (2018). Non-O157 Shiga toxin-producing Escherichia coli-A poorly appreciated enteric pathogen: Systematic review. International Journal of Infectious Diseases:UID: official publication of the Interational Society for Infectious Diseases,76: 82–87.

57. Valilis, E., A. Ramsey, S. Sidiq, and H. DuPont. Non-O157 Shiga toxin-producing Escherichia coli – A poorly appreciated enteric pathogen: Systematic review. (2018). International Journal of Infectious Diseases, 76: 82–87.

58. Wang, C., C. Hsu, H. Huang, and B. B. Yang. (2016). Recent advances in food processing using high hydrostatic pressure technology. Critical Reviews in Food Science and Nutrition 56,(4):527–540.

59. Xu, Y., A. Scales, K. Jordan, C. Kim, and E. Sismour. (2016). Starch nanocomposite films incorporating grape pomace extract and cellulose nanocrystal. Journal of Applied Polymer Science, 134(6): 44438.

60. Yaldagard, M., S.A. Mortazavi, and F. Tabatabaie. (2008). The principles of ultra high pressure technology and its application in food processing/preservation: A review of microbiological and quality aspects. African Journal of Biotechnology, 7(16): 2739–2767.

61. Yordanov, D., and G.V. Angelova. (2010). High Pressure Processing for Food Preserving. Biotechnology & Biotechnological Equipment, 24(3): 1940–1945.

62. Zhu, M., M. Du, J. Cordray, and D.U. Ahn, (2005). Control of Listeria monocytogenes contamination in ready to eat meat products. Comprehensive Reviews in Food Science and Food Safety,4:34–42.

63. Zielinski, A.A.F., D.M. Zardo, A. Alberti, D.G. Bortolini, L. Benvenutti, I.M. Demiate, and A. Nogueira. (2019). Effect of cryoconcentration process on phenolic compounds and anxtioxidant activity in apple juice. Journal of the Science of Food and Agriculture, 99(6): 2786–2792.

